# Targeting tumor-intrinsic TAK1 triggers anti-tumor immunity and sensitizes pancreatic cancer to checkpoint blockade

**DOI:** 10.1101/2025.10.08.681226

**Authors:** Sapana P. Bansod, Timothy Hung-Po Chen, Vikas K. Somani, Lin Li, Suneeta Modekurty, Brett L. Knolhoff, Ryan C. Fields, Marianna B. Ruzinova, David G. DeNardo, Kian-Huat Lim

**Author notes:** Corresponding author: Kian-Huat Lim, Washington University School of Medicine 660 South Euclid Avenue, Campus Box 8069 Saint Louis, MO 63110. The authors have declared no conflicts of interest. All data, analytic methods, and study materials will be made available to other researchers upon request to the corresponding author. All genomic data has been deposited at Gene Expression Omnibus.

## Abstract

**Background and Aims:** Targeting the Transforming Growth Factor-β (TGF-β) pathway to reverse the immunologically “cold” tumor microenvironment (TME) of pancreatic ductal adenocarcinoma (PDAC) remains clinically unsuccessful, warranting novel therapeutic strategies.

**Methods:** We developed a novel tumor-CD8 T cell co-culture to interrogate the TGF-β signaling pathways that promotes T cell-mediated cytotoxicity. We performed multiplex immunohistochemistry (mIHC) on human PDAC samples to correlate cell-type specific TGF-β pathway activation and CD8 T cell abundance. We employed specific pathway inhibitor and newly generated genetically-engineered mouse models (GEMMs) and confirmed our findings using single-cell RNA sequencing, flow cytometry and mIHC. We performed proteomics and various in vitro assays to establish the molecular mechanisms.

**Results:** We identify TGF-β-activated kinase 1 (TAK1 or MAP3K7) as an aberrantly activated kinase in human and mouse PDAC tissues that is associated with T cell dysfunction. Pharmacological inhibition of TAK1 with Takinib, or genetic deletion of *MAP3K7* in autochthonous *p48-Cre;TP53^flox/flox^;LSL-KRAS^G12D^*GEMM, enhances intratumoral CD4^+^ and CD8^+^ effector T cell infiltration and renders immune checkpoint blockade (ICB) effective. Mechanistically, TAK1 inhibition induces DNA damage and cytoplasmic DNA leakage, which activates the cyclic GMP-AMP synthase–Stimulator of Interferon Genes (cGAS–STING) DNA sensing pathway, triggering inflammatory responses that promote adaptive immune cell infiltration. At the molecular level, TAK1 phosphorylates Ephrin Receptor A2 (EphA2) at Serine 897, which in turn phosphorylates RAD51 at Tyrosine 315, a key DNA repair protein involved in homologous recombination.

**Conclusions:** We uncover TAK1 as a critical mediator in maintaining genomic integrity and highlights its potential as a therapeutic target to induce an inflamed TME that sensitizes PDAC to ICB.

## INTRODUCTION

Pancreatic ductal adenocarcinoma (PDAC) is lethal malignancy characterized by a chronically inflamed and desmoplastic tumor microenvironment (TME), which drives chemoresistance and immune evasion resulting in poor responsiveness to immune checkpoint blockade (ICB)^1^. Inflammatory signaling such as the IL-1R/TLR-NF-κB and TGF-β−SMAD2/3/4 cascades not only propel neoplastic progression but also reprogram the CAFs and immune infiltrates to foster a desmoplastic TME^2^, supporting the rationale to combine inhibitors of these pathways with ICB.

TGF-β signaling operates through both SMAD-dependent and SMAD-independent (non-SMAD) pathways. In the canonical SMAD pathway, receptor-regulated SMADs (SMAD2/3) are phosphorylated by the activated TGF-β receptor complex, then partner with SMAD4 to translocate to the nucleus and regulate gene transcription. Non-SMAD pathways involve activation of parallel signaling cascades, such as the MAPK, PI3K/AKT, TAK1 and Rho-like GTPases, that modulate cellular responses independently or synergistically with SMAD signaling^3^. In this study, we identify TAK1 (Transforming Growth Factor β-Activated Kinase 1, also known as MAP3K7) as a key mediator that causes T cell dysfunction in PDAC. We found that TAK1 inhibition induces DNA damage and subsequently cGAS-STING pathway activation, resulting in an inflammatory milieu that is T cell -permissive and renders ICB effective.

## MATERIALS AND METHODS (Please see Supplementary Data for more information)

### DATA TRANSPARENCY STATEMENT

Bulk RNAseq data is deposited in GEO with accession number GSE293275 and GSE293275 and and Murine scRNAseq is deposited in GEO with accession number GSE297798 and GSE297618. All data, analytic methods, and study materials will be made available to other researchers upon request to the corresponding author.

## RESULTS

### TAK1 is activated in the epithelial compartment of PDAC and is associated with T cell dysfunction

The advent of single-cell(sc) RNAseq have enabled analysis of dysregulated signaling pathways within each cellular compartments of human tumors. To this end, we pooled and re-analyzed scRNA-seq datasets of normal pancreatic (n=14) and PDAC tissues (n=40) published in three different studies^4–6^(**Fig. 1A**). Focused gene set enrichment analysis (GSEA) on the epithelial compartment showed enrichment of RAS, ERK pathways, as expected, and also the TNF-NF-κb, TGF-β and interleukins inflammatory pathways (**Fig. 1B, 1C, Suppl. Fig. 1A**), which have all been implicated in immune evasion^7^. Of these, we focused on TGF-β with the goal of identifying novel therapeutic nodes within this pathway that can potentiate ICB.

**Figure 1.**
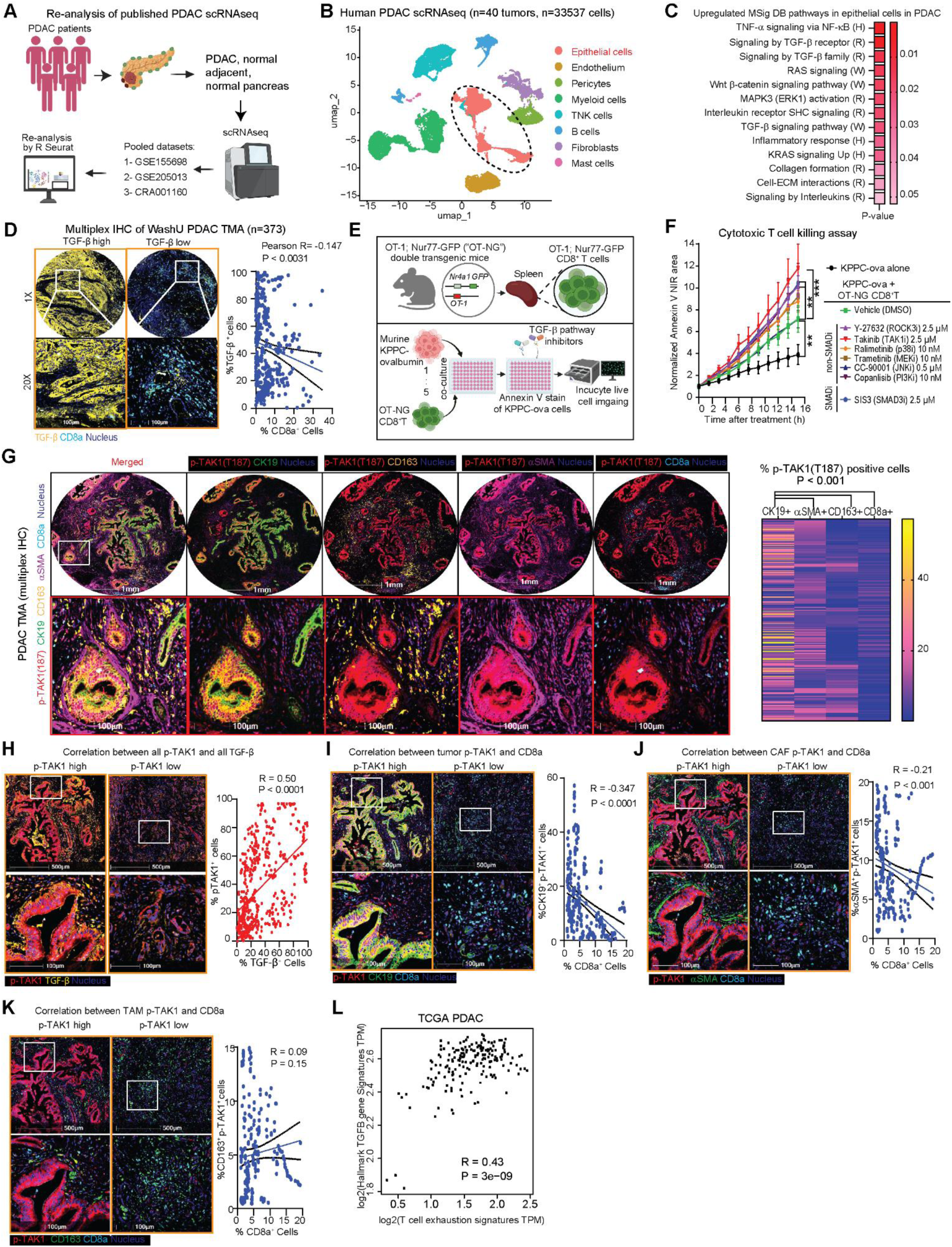
TAK1 is activated in the epithelial compartment of PDAC and is associated with T cell dysfunction. **(A)** Schematic of human PDAC scRNAseq from 3 datasets, merged and analyzed with R Seurat. **(B)** UMAP plot scRNAseq data from human PDAC tumors showing 8 different cell clusters. **(C)** Heatmap of epithelial cells of PDAC, GSEA datasets: Hallmark (H), Reactome (R), and WikiPathways (W). **(D)** Representative multiplex IHC images of a human PDAC TMA with high and low TGF-β staining (yellow) and a correlation plot with CD8a (magenta). **(E)** Schematic showing isolation and co-culture of OT-NG CD8+ T cells from the spleens of double-transgenic OT-1; Nur77-GFP (OT-NG) mice and KPPC-ova cells. **(F)** Normalized Annexin V NIR area of TGF-β pathway inhibitors as indicated time (N=5/group). **(G)** Representative multiplex IHC images and heatmap of % colocalization of phospho-TAK1 (T187, red) with each cell-specific marker: CK19+ cells (green), CD163+ cells (orange), αSMA+ cells (purple), and CD8a+ cells (magenta) in the PDAC TMA. **(H)** Representative IHC images of PDAC TMA with high and low phospho-TAK1 (T187, red) and correlation plot with TGF-β (yellow). **(I)** Representative IHC images of PDAC TMA samples with high and low phospho-TAK1 (T187, red) CK19+ cells (green), **(J)** αSMA+ cells (green), **(K)** CD163+ cells respectively and its correlation plots with CD8a+ cells (magenta). **(L)** Correlation plots of Hallmark TGF-β vs. T cell exhaustion signatures in the TCGA PDAC database from GEPIA 2. (*P < 0.05, **P < 0.01, ***P < 0.001; scale bars 500 or 100 µm).

We performed multiplex immunohistochemistry (mIHC) on a human PDAC tissue microarray (TMA) consisting of 373 different surgical PDAC samples. Across these samples we observed a negative correlation between global TGF-β staining intensity and CD8a T cell abundance (**Fig 1D**). Notably, TGF-β staining appeared most prominent in PDAC cells, corroborating the scRNAseq data showing upregulation of TGF-β signature (**Fig. 1C**). To identify the downstream signaling that plays a role in immune evasion, we developed an in vitro tumor and antigen-specific T cell co-culture assay. We crossed C57BL/6J transgenic OT-1 mice, which T cells express MHC I-restricted ovalbumin 257-264^8^, and C57BL/6J Nur-77-GFP reporter mice, whose T cells express green fluorescent protein (GFP) upon stimulation of T cell receptor^9^, to obtain double OT-1; Nur-77-GFP (OT-NG) transgenic mice. We also engineered a murine PDAC cell line derived from a *p48-Cre;Trp53^flox/flox^;LSL-Kras^G12D^*PDAC mouse to stably express ovalbumin (KPPC-ova, **Fig. 1E**). When co-cultured, the OT-NG CD8^+^ T cells were able to kill KPPC-ova cells, as measured by Annexin V NIR-dye using Incucyte® (**Suppl. Fig 1B**). We then treated this co-culture with different TGF-β pathway inhibitors at concentrations that were not lethal to KPPC-ova cells, as determined by YOYO-1 fluorescence (**Suppl. Fig. 1C**). While adding OT-NG CD8^+^ T cells resulted in increased death of KPPC-ova cells, as measured by Annexin V NIR signal, adding TAK1, PI3K and SMAD3 inhibitors into the co-culture further augmented the killing effect (**Fig. 1F, Suppl. Fig. 1D**) without diminishing GFP signal in the co-culture, indicating no harm to CD8^+^ T cell activity (**Suppl. Fig. 1E**). We chose to focus on TAK1 due to its relatively underexplored role in the immune TME of PDAC.

To understand the role of TAK1 signaling in human PDAC, we first performed mIHC on PDAC TMA. using a phospho-TAK1 (T187) antibody which identifies TAK1 activation, along with cell-type-specific markers. We found that phospho-TAK1 (T187) is detected more prominently in PDAC (CK19, 45.9%) cells, followed by CAFs (αSMA, 27.6%), macrophages (CD163, 15.5%) and CD8^+^ T cells (10.8%, **Fig. 1G**). Notably, phospho-TAK1 positively correlated positively with TGF-β staining intensities across the PDAC samples (**Fig. 1H**). The abundance of phospho-TAK1 staining in PDAC cells (**Fig. 1I**), and to a lesser extent, CAFs (**Fig. 1J**), negatively correlated with CD8^+^ T cell abundance. Presence of phospho-TAK1 in macrophages did not correlate with CD8^+^ T cell abundance (**Fig. 1K**). Analysis of the PDAC database TCGA, which reports bulk RNAseq data, showed higher expression of *Map3k7* expression in PDAC compared to normal pancreatic tissue (**Suppl. Fig. 1F**), positive correlation between *Tgfb1* and *Map3k7* expression in PDAC (**Supple. Fig 1G**). Hallmark *Tgfb* signature and expression of *Tgfb1* or *Map3k7* also positively correlated with T cell exhaustion signature^10^ (**Fig. 1L, Suppl. Fig 1H**). Higher expression of *Map3k7* is also associated with poorer disease-free survival (DFS) and overall survival (OS, **Suppl. Fig. 1I**). Together, these data led us to hypothesize that tumor-intrinsic TAK1 activation leads to T cell dysfunction.

### Suppression of tumor-intrinsic *Map3k7* reprograms the immune TME

Besides fully developed PDAC cells, activated phospho-TAK1 (T187) staining is present in early and late PanIN in both human (**Fig. 2A**) and autochthonous KPPC mice (**Fig. 2B**), suggesting a role of TAK1 activity in PDAC development. To understand the role of TAK1 in PDAC tumorigenesis, we silenced *Map3k7* in a KPPC cell by two different small hairpin (sh) RNAs, or a scrambled RNA sequence as control (scram.), and injected them subcutaneously into C57BL/6J or nude mice. We found *Map3k7*-silenced KPPC cells were impaired in forming tumors in C57BL/6J, but not in nude mice (**Fig. 2C**), suggesting a role of an intact immune system in suppressing the tumorigenesis of *Map3k7*-silenced KPPC tumors. To show this, we inoculated KPPC cells into the pancreas of Nur77-GFP reporter mice the OT-1 transgene and performed immunofluorescence (IF) of the orthotopic tumors two weeks later (**Fig. 2D**). We found significantly more GFP^+^ CD8^+^ T cells in *Map3k7*-silenced tumors, as compared to control tumors (**Fig. 2E**). Next, we performed FACS analysis on control and *Map3k7*-silenced orthotopic tumors two weeks after inoculation. Consistent with subcutaneous model, *Map3k7*-silenced tumors were reduced in weight (**Fig. 2F**), infiltrated with more total and activated effector (CD44^high^Ki-67^+^) CD4^+^ and CD8^+^ T cells, and less tumor associated macrophages (TAMs), which were more M1-polarized (**Fig. 2G**). This data provides robust evidence that tumor-intrinsic TAK1 signaling plays a suppressive role in adaptive T cell immunity. To understand the underlying mechanism, we performed bulk-RNAseq *Map3k7*-silenced KPPC cells (**Fig. 2H**). Surprisingly, pathway analysis showed enrichment of inflammatory signatures including the TNF, IL-6, IFNα and IFNγ, TLR and NLR pathways. However, TGF-β signature was downregulated, consistent with the known role of TAK1 as a downstream driver of TGF-β signaling cascade (**Fig. 2I**, **Fig. 2J**). We independently confirmed the RNA-seq results by qPCR that *Map3k7*-silenced cells expressed significantly higher *Tnf, Il6, Il1a, Il1b, Nlrp3* and less *Tgfb1* (**Suppl. Fig. 2A**). Interestingly, *Map3k7*-silenced cells displayed significantly downregulated DNA repair and G2M checkpoint signatures (**Fig. 2I**, **Fig. 2J, Suppl. Fig. 2B, 2C**), suggesting a previously unappreciated role of TAK1 signaling in DNA repair and cell cycle progression.

**Figure 2.**
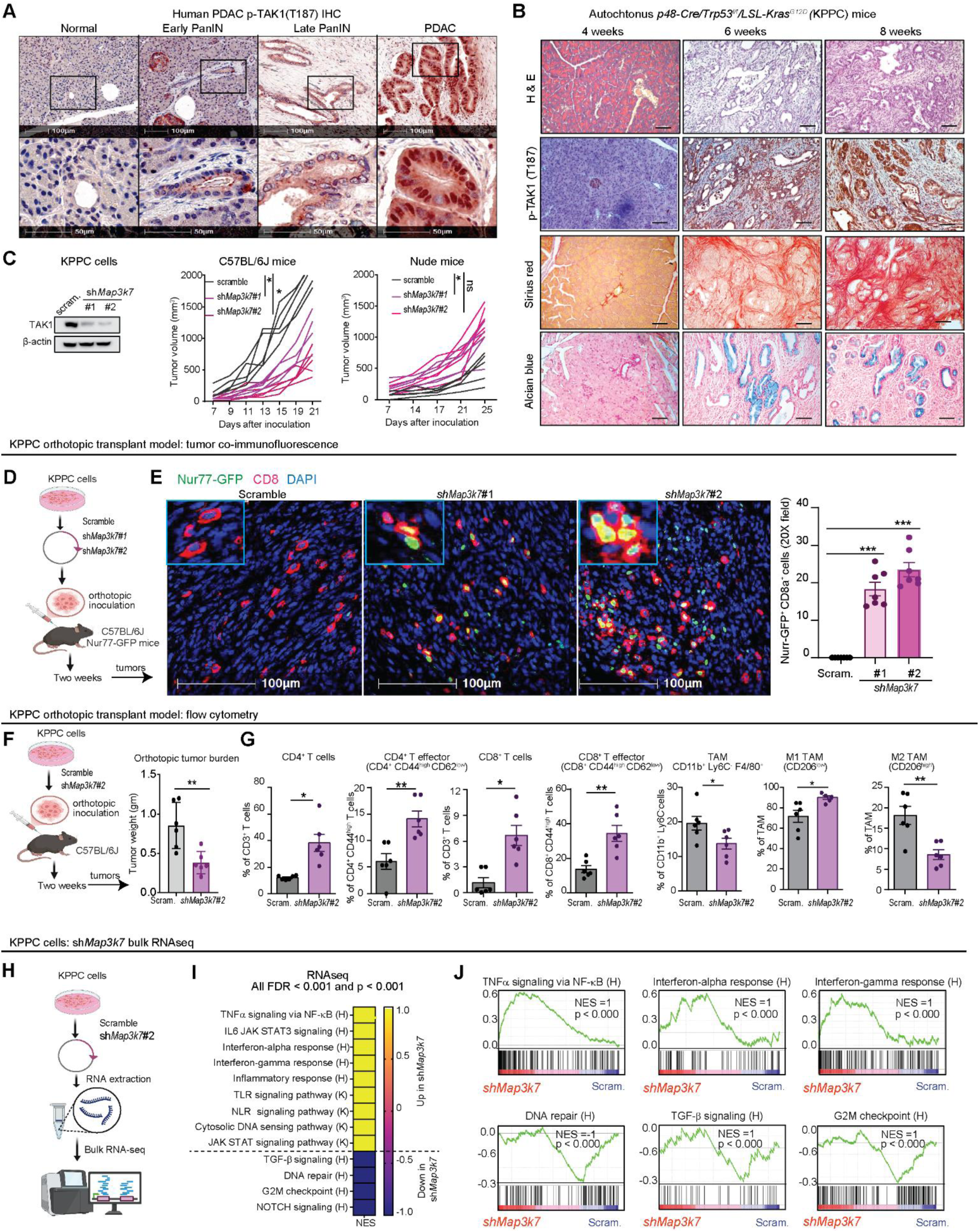
Suppression of tumor-intrinsic *Map3k7* reprograms the immune TME. **(A)** Representative IHC images of phospho-TAK1 (T187) staining in human normal pancreatic tissues, early PanIN, late PanIN, and PDAC. **(B)** Representative images of H&E, phospho-TAK1 IHC, Sirius red, and Alcian blue staining from KPPC mice at different ages. **(C)** Western blots of TAK1 in KPPC cells stably expressing two different shRNAs against *Map3k7* and their growth kinetics in C57BL/6J and nude mice. **(D)** Experimental schematic of scram. or sh*Map3k7* KPPC cells inoculated orthotopically into Nur77-GFP reporter mice (N = 7 mice/arm). **(E)** Representative co-IF images and quantification of Nur77-GFP+ (green) and CD8+ (red) T cells in scram. or sh*Map3k7* KPPC tumors. **(F)** Experimental schematic and weight of tumors from C57BL/6 mice orthotopically inoculated with scram. or sh*Map3k7* KPPC cells (N=6 mice/arm). **(G)** FACS quantification of intratumoral T and myeloid cells panel in scram. or sh*Map3k7* KPPC tumors. **(H)** Schematic showing RNAseq, **(I)** Heatmap and **(J)** GSEA plots of selected signatures in sh*Map3k7* KPPC cells. Kyoto Encyclopedia for Genes and Genomes (K), and Hallmark (H) database from MSigDB. (\**P* < 0.05, \*\**P* < 0.01, \*\*\**P* < 0.001, ns not significant; scale bars 50 and 100 µM).

### Conditional *Map3k7*-deleted KPPC mice produce PDAC with enhanced T cell infiltration

To robustly study the role of tumor-intrinsic TAK1 in development of PDAC and the native TME, we obtained a *Map3k7^f/wt^* B6;129S7 mouse in which exon 1 of the *Map3k7* gene was flanked by the loxP sequences^11^, and backcrossed it nine times to C57BL/6J mice before crossing it with the C57BL/6J KPPC parental strains to generate conditional heterozygous (*Map3k7^f/wt^*) and homozygous (*Map3k7^f/f^*) *Map3k7*-knockout KPPC mice in pure C57BL/6J background. Interestingly, these two mouse strains still developed PDAC but at vastly different kinetics. IHC staining of the pancreata from these mice showed complete loss of phospho-TAK1 staining in the epithelia of *Map3k7^f/wt^*and *Map3k7^f/f^* KPPC tumors, respectively, confirming correct genotyping (**Fig. 3A, Suppl. Fig. 3A**). We performed multiple analyses on the pancreata of age-matched mice (**Fig. 3B**). Compared to wild-type (WT), heterozygous *Map3k7^f/wt^* KPPC mice had a delayed acinar-to-ductal transition, neoplastic progression from PanIN to PDAC and stromal expansion by Sirius red staining (**Fig. 3C**). The *Map3k7^f/wt^* KPPC mice survived significantly longer than WT (**Fig. 3D**), and when euthanized, had lower PDAC tumor burden (**Fig. 3E**). On the contrary, homozygous *Map3k7^f/f^* KPPC mice underwent acinar-to-ductal metaplasia, formed PanIN and had stromal expansion as early as 4 weeks after birth, when the pancreata of WT KPPC mice were completely normal (**Suppl. Fig 3B, Suppl. Fig. 3C**). However, at 8 weeks of age when the pancreata of WT mice were mostly occupied with PDAC, the pancreata of *Map3k7^f/f^* KPPC mice were almost completely necrotic with scant foci of PDAC (**Suppl. Fig 3B**). The early transition to PanIN and depletion of pancreas probably contributed to the shorter survival of *Map3k7^f/f^* KPPC mice (**Suppl. Fig 3D**), which when euthanized had minimal pancreas weight (**Suppl. Fig. 3E**).

**Figure 3.**
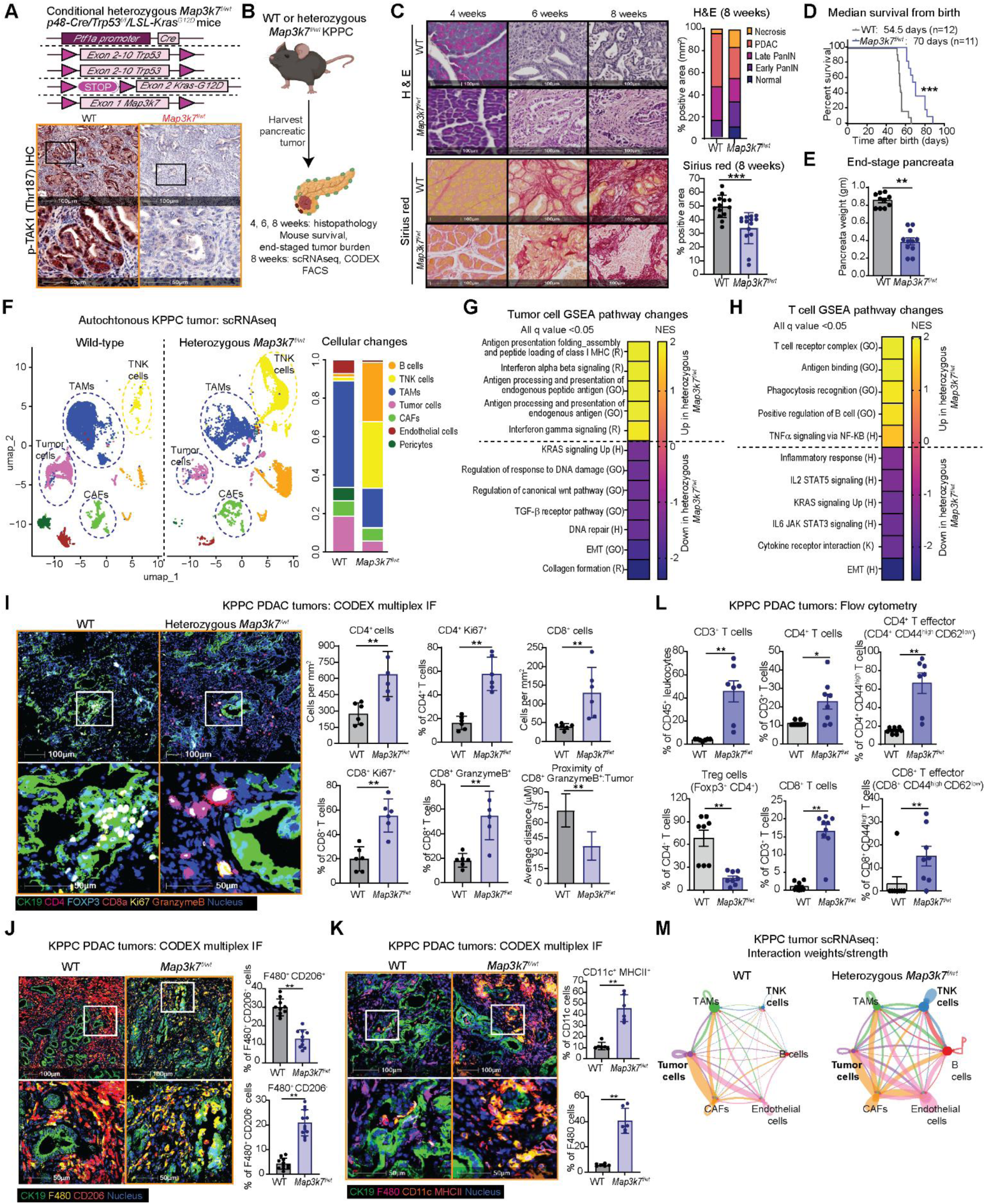
Conditional *Map3k7*-deleted KPPC mice produce PDAC with enhanced T cell infiltration. **(A)** Conditional strategy to delete one *Map3k7* allele in KPPC mice and IHC of phospho-TAK1 (T187) in *Map3k7^f/wt^* KPPC tumors. **(B)** Schematic diagram showing experiments performed on wildtype (WT) and *Map3k7^f/wt^* KPPC mice. **(C)** Representative images and quantification of H&E and Sirius red staining of 4-, 6-, and 8-weeks old WT and *Map3k7^f/wt^* KPPC tumors. **(D)** Kaplan-Meier survival plot **(E)** Tumor burden of the pancreata of WT and *Map3k7^f/wt^*KPPC mice at humane endpoints. **(F)** UMAP plot of scRNAseq of WT and *Map3k7^f/wt^*KPPC tumors showing cell subsets (N=4 tumors/arm). Heatmaps showing GSEA signatures in *Map3k7^f/wt^* **(G)** tumor or **(H)** TNK cells based on scRNAseq. Hallmark (H), Gene Ontology (GO), Reactome (R) and Kyoto Encyclopedia for Genes and Genomes (K), database from MSigDB. Representative CODEX-based multiplex IF images and quantification of **(I)** T cells panel and proximity analysis; **(J)** F480^+^ CD206^+^ and F480^+^ CD206^-^ TAMs; and (**K)** CD11c^+^ MHC II^+^ dendritic cells and F480^+^ MHC II TAMs. **(L)** FACS quantification of intratumoral T cells panel of WT and *Map3k7^f/wt^* KPPC tumors (N=8/arm). (**M)** Differential interaction weight/strength between TNK and major cell types from WT and *Map3k7^f/wt^* KPPC tumors by CellChat analysis. (\**P* < 0.05, \*\**P* < 0.01, \*\*\**P* < 0.001; scale bars 50 and 100 µM).

To gain a depth understanding on the TME, we performed scRNAseq analysis on PDAC tumors from WT, heterozygous *Map3k7^f/wt^* and homozygous *Map3k7^f/f^*KPPC mice at 8 weeks of age. The homozygous *Map3k7^f/f^* KPPC tumors contained significantly fewer total cells, which did not permit robust pathway analyses (**Suppl. Fig. 3F, 3G**). Uniform manifold approximation and projection (UMAP) clustering showed that the heterozygous *Map3k7^f/wt^* KPPC tumors contained more T and natural killer (NK) cells, and less PDAC and TAMs (**Fig. 3F, Suppl. Fig. 3H**). GSEA of PDAC cells showed upregulation of IFN and antigen-presentation, as well as downregulation of TGF-β and DNA repair signatures in of *Map3k7^f/wt^* KPPC tumors (**Fig. 3G**), which recapitulate the RNAseq (**Fig. 2I**). The TNK cells in *Map3k7^f/wt^* KPPC tumors were enriched for Cd8, had higher expression of *Gzma, Tnf* and *Ifng* (**Suppl. Fig 3I**), and displayed upregulated T cell receptor, antigen binding, phagocytosis, and TNF signatures (**Fig. 3H**), indicative of an activated state. We confirm the scRNAseq data by multiplex IF using CO-Detection by indexing (CODEX®) technique. Compared to WT, *Map3k7^f/wt^* tumors had lower αSMA staining, consistent with lower Sirius red and lower TGF-β staining, which we also observed in *Map3k7*-silenced cells (**Suppl. Fig. 3J**), suggesting a positive role of TAK1 in TGF-β expression in PDAC cells. *Map3k7^f/wt^* tumors had more total and proliferating intratumoral CD4^+^, CD8^+^ T cells, cytotoxic (granzyme B^+^) CD8^+^T cells (**Fig. 3I**), less CD206^+^ (an M2 marker) TAMs (**Fig. 3J**) and more MHC II^+^ F480^+^ TAMs and CD11c^+^ dendritic cells (**Fig. 3K**). As independent confirmation, we performed flow cytometry on 8-week-old WT and *Map3k7^f/wt^* KPPC tumors. Again, *Map3k7^f/wt^* tumors had an increased abundance of intratumoral CD3^+^ T cells, total and effector CD4^+^ and CD8^+^ T cells and a decreased proportion of CD4^+^

Foxp3^+^ regulatory T cells (**Fig. 3L**), as well as much lower abundance of granulocytes, monocytes, M2-like TAMs which were skewed towards M1-polarization (**Suppl. Fig 3K**). These data showed that suppression of tumor TAK1 reprograms the immune TME to favor an adaptive anti-tumor T cell response. Supporting this notion, CellChat analysis of the scRNAseq data nominated increased interactions of various cell types, including PDAC cells, with TNK cells in *Map3k7^f/wt^* tumors (**Fig. 3M**). Focused analyses of interaction between PDAC and TNK cells suggested an increased antigen presentation by MHC molecules from tumor to CD8^+^ T cells (**Suppl. Fig. 3L**). This data led us to hypothesize that suppression of TAK1 led to tumor-intrinsic changes that elicit T cell response.

### TAK1 suppression results in DNA damage and cGAS-STING activation

TAK1 is a crucial kinase that transmit signaling downstream of multiple inflammatory receptors including the IL-1/Toll-like receptors, TNF and the non-SMAD aspect of TGF-β receptors^12^. Surprisingly, several inflammatory cytokines including *Ifna, Ifnb, Ifng, Il1, Il6, Tnf* and their signatures were paradoxically upregulated in *Map3k7*-silenced and *Map3k7^f/wt^* KPPC cells (**Suppl. Fig. 4A**). Additionally, *Map3k7*-silenced KPPC cells had downregulated TGF-β, DNA repair signatures but elevated cytosolic DNA sensing signatures (**Fig. 2J**, **Suppl. Fig. 4B**). These findings led us to hypothesize that TAK1 may be crucial for DNA damage repair and when inhibited. To study the molecular mechanism, we performed reverse phase protein array (RPPA) analysis on scramble-control and two different *Map3k7*-silenced KPPC cells. In *Map3k7*-silenced cells, several markers such as phospho-EGFR(Y1173), Snail, phospho-c-Jun(S72), phospho-EphA2 (S897) and β-catenin were downregulated. On the other hand, several markers were upregulated include phospho-YAP1(S127), STING, IRF3, phospho-p65(S536), ATM and STAT3 (**Fig. 4A, Suppl. Fig. 4C**). In support, expressions of *Irf1, Irf3, Irf7, Tbk1, Stat1* and *Cgas* were all upregulated in *Map3k7*-silenced KPPC cells (**Suppl. Fig. 4D**). These findings demonstrate that TAK1 inhibition triggers cGAS-STING cascade. Indeed, by western blots *MAPK37*-silenced KPPC cells displayed upregulated cGAS-STING markers (**Fig. 4B**), and qPCR also showed upregulated expressions of *Irf1, Irf3, Cgas, Sting, Ifnr1, Ifnr2, Ifna, Ifnb* and *Ifng* (**Fig. 4C**). In support of these results, treatment of KPPC cells with Takinib, a specific TAK1 inhibitor^13^, also increased phospho-STING, cGAS and phopho-STAT1 (**Fig. 4D**). Additionally, multiplex IHC of *Map3k7^f/wt^* KPPC tumors showed increased phospho-TBK1 and STING staining in CK19^+^ PDAC cells (**Fig. 4E**).

**Figure 4.**
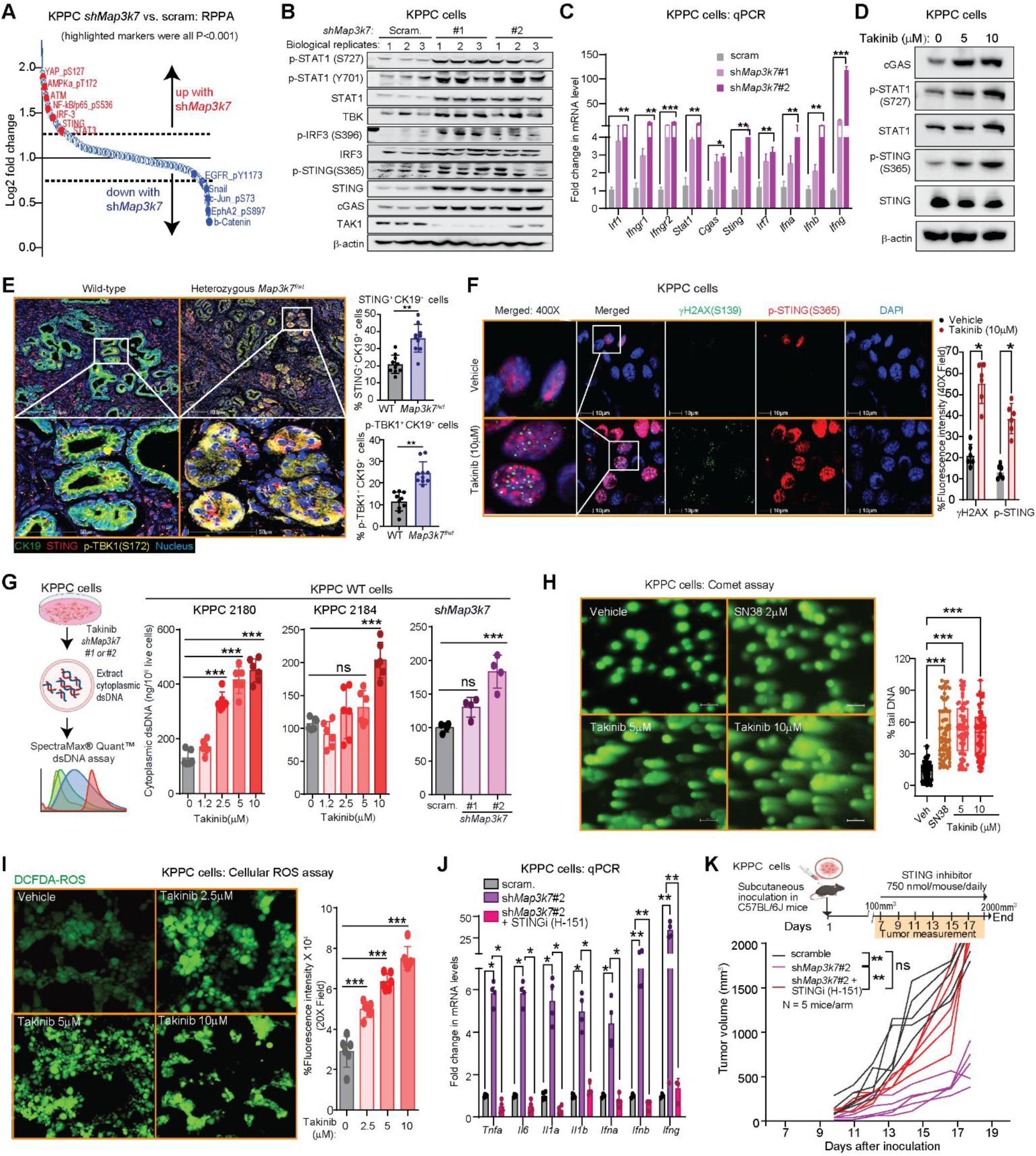
TAK1 suppression results in DNA damage and cGAS-STING activation. **(A)** RPPA showing significant up- or downregulated markers in *shMap3k7* when compared to scram. KPPC cells. **(B)** Western blots and **(C)** qPCR of the indicated proteins or genes in *shMap3k7* and scram. KPPC cells. **(D)** Western blot of indicated proteins in KPPC cells treated with Takinib overnight. **(E)** Representative multiplex IF images and quantification of CK19 (green), STING (red) and phospho-TBK1(S172) (yellow) in WT and *Map3k7^f/wt^*KPPC tumors. **(F)** Representative IF images of ϒH2AX(S139) and phospho-STING(S365) staining in KPPC cells treated as indicated overnight. **(G)** Experimental schematic and quantification of cytoplasmic dsDNA in KPPC cells treated as indicated overnight in two different WT, or in *shMap3k7* KPPC line. **(H)** Representative images and quantification of Comet assay in KPPC cells treated as indicated for overnight. **(I)** Representative images and quantification of ROS by DCFDA in KPPC cells treated as indicated for overnight. **(J)** qPCR of inflammatory genes in *shMap3k7* vs. scram. and treated as indicated for overnight. (**K)** Experimental schematic and tumor kinetics of scram. and sh*Map3k7* KPPC tumors in C57BL/6J mice and treated with STING inhibitor (N=5 mice/arm). (\**P* < 0.05, \*\**P* < 0.01, \*\*\**P* < 0.001; scale bars 50 and 100 µM).

Because the cGAS-STING cascade is triggered when cells sense cytosolic DNA, we investigated whether TAK1 inhibition leads to DNA damage, and thus DNA leakage into the cytoplasm. Indeed, Takinib treatment resulted in increased γH2AX (S139), a marker of DNA double-strand breaks (DSBs) in addition to increased phospho-STING (**Fig. 4F**). Additionally, suppression of TAK1 by shRNA, heterozygous deletion or Takinib increased cytosolic DNA abundance, as measured by SpectraMax® Quant^TM^ fluorometric assay (**Fig. 4G, Suppl. Fig. 4E**). Furthermore, Takinib treatment increased DNA tails of individual cells, as also seen with topoisomerase I inhibitor SN38, by alkaline comet assay, an established method to detect DNA strand breaks^14^ (**Fig. 4H**). Besides inflammation, cGAS-STING activation also leads to accumulation of reactive oxygen species (ROS), which we observed in Takinib-treated KPPC cells using the cell permeable 2’,7’-dichlorodihydrofluorescein diacetate (DCFDA) fluorescent probe^15^ (**Fig. 4I**).

Activation of the cGAS-STING pathway triggers inflammatory responses that can potentially transform immunologically “cold” tumors into “hot” tumors, thereby enabling T cell responses^16^. In KPPC cells, silencing of *Map3k7* upregulated expression of inflammatory cytokines including *Ifna, Ifnb, Ifng, Tnf, Il1a, Il1b* and *Il6*, as well as phospho-STAT1, but these could be blocked by STING inhibitor H-151 (**Fig. 4J, Suppl. Fig. 4F**). Importantly, the suppressed tumorigenesis of KPPC cells in C57BL/6J mice after silencing of *Map3k7* was abrogated when mice were treated with H-151 (**Fig. 4K**). Collectively, these studies demonstrate that TAK1 inhibition results in DNA damage and sensing of the cytosolic DNA by the cGAS-STING cascade.

### TAK1 phosphorylates EphA2 and RAD51 to maintain DNA integrity

Following DNA damage, TAK1 is recruited in the cytoplasm by the ATM/NEMO/RIP1 complex to promote cell survival while allowing DNA repair to occur^17^. However, TAK1 has not been reported to directly involve in DNA damage repair, nor is it known to have a role in the nucleus. In human and mouse PDAC, TAK1 antibody stains predominantly the cytoplasm by IHC, as reported^18^. When activated, TAK1 autophosphorylates Thr-184 and -187 within its activation loop to unleash its kinase activity^19^. Interestingly, we found that phospho-TAK1 (T187) staining is present in both the cytoplasm and the nucleus of human and wild-type KPPC PDAC cells (**Fig. 5A**) and is markedly diminished or absent in *Map3k7^f/wt^* and *Map3k7^f/f^* PDAC cells, respectively, attesting to the specificity of the antibody. Cultured PDAC cells also showed prominent phospho-TAK1 staining in the nucleus, but the intensity was diminished when the cells were treated with Takinib (**Fig. 5B**). These data suggest that TAK1 may have a function in the nucleus or DNA repair, but the molecular mechanism is unclear.

**Figure 5.**
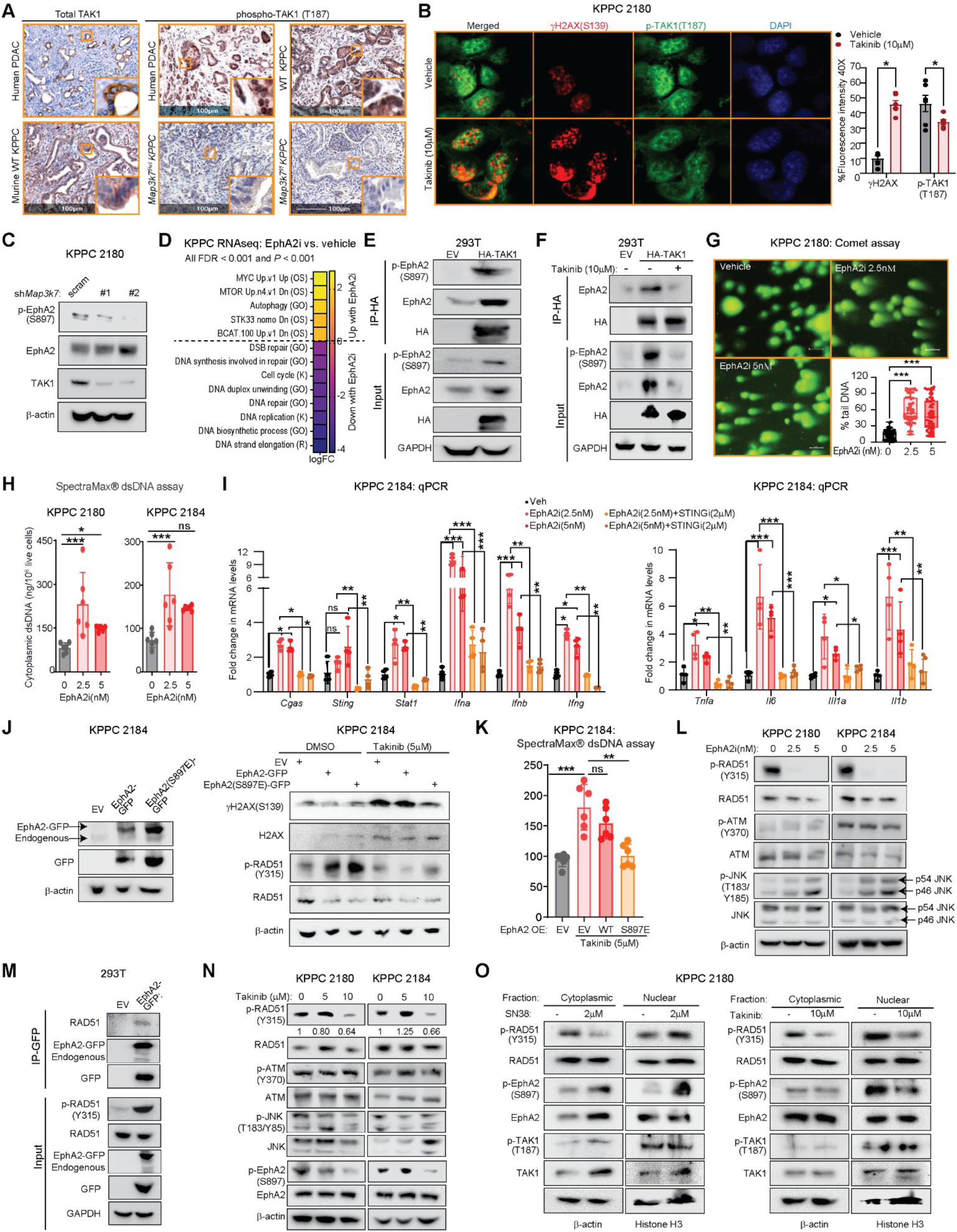
TAK1 phosphorylates EphA2 and RAD51 to maintain DNA integrity. **(A)** Representative IHC images of TAK1 and phospho-TAK1 (T187) staining in human and murine PDAC from WT, *Map3k7^f/wt^* and *Map3k7^f/f^* KPPC mice. **(B)** Representative IF images and quantification of ϒH2AX(S139) and phospho-TAK1 (T187) staining treated as indicated overnight. **(C)** Western blots of *shMap3k7* KPPC cells. **(D)** Heatmap showing GSEA signatures in KPPC cells treated EphA2 inhibitor (ALW-II-41-27, 2.5nM) for overnight. Kyoto Encyclopedia for Genes and Genomes (K), Reactome (R), Oncogenic signature (OS) and Gene Ontology (GO) database from MSigDB. **(E, F)** IP-western blots 293T cells treated with Takinib overnight. **(G)** Representative images and quantification of Comet assay and **(H)** Quantification of cytoplasmic dsDNA in WT KPPC cell lines treated with vehicle or EphA2 inhibitor for overnight. **(I)** qPCR showing cGAS-STING genes in KPPC cells treated as indicated overnight. **(J)** Western blots and **(K)** Quantification of cytoplasmic dsDNA in KPPC cells stably expressing an empty vector (EV), EphA2(WT)-GFP or EphA2(S897E)-GFP treated with Takinib (5μM) for overnight. **(L)** Western blot of EphA2 inhibitor treated WT KPPC cell lines overnight. **(M**) IP-western blots in 293T cells ectopically-expressed EphA2-GFP. (**N)** Western blot of indicated markers treated as indicated overnight. **(O)** Western blots of cytoplasmic or nuclear proteins of KPPC cells treated as indicated for overnight. (\**P* < 0.05, \*\**P* < 0.01, \*\*\**P* < 0.001, ns: not significant; scale bars 50 or 100 µM).

Our RPPA data showed that in *Map3k7*-silenced KPPC cells, phosphorylation of Ephrin type-A receptor 2 receptor tyrosine kinase (EphA2 RTK) at S897 is the most significantly downregulated marker (**Fig. 4A**), which we confirmed by western blots (**Fig. 5C**). Phospho-EphA2(S897) has been shown to be upregulated following irradiation to promote survival in multiple cancer types^20^. Interestingly, a chemical genetic kinase screen identified EphA2 as a potential substrate for TAK1 in PDAC cell lines^21^. We therefore hypothesize that TAK1 may engage EphA2 as a substrate in DNA repair. First, we performed RNAseq on KPPC cells treated with vehicle or ALW-II-41-27^22^, a potent EphA2 inhibitor (EphA2i). Interestingly, EphA2-treated cells displayed downregulation of multiple DNA repair signatures, supporting a mechanistic link between TAK1 and EpHA2 (**Fig. 5D**). By co-immunoprecipitation (co-IP), TAK1 binds and phosphorylates EphA2 (**Fig. 5E**), and the interaction is diminished by Takinib (**Fig. 5F**). Additionally, EphA2i at 2.5nM induced DNA strand breaks (**Fig. 5G**), cytoplastic DNA (**Fig. 5H**), and expression of *Cgas, Sting* and other inflammatory genes in a STING-dependent manner (**Fig. 5I**). These data show that EphA2 is crucial in maintaining DNA integrity in PDAC cells. To demonstrate the relationship between TAK1 and EphA2, we generated KPPC cells stably expressing empty vector (EV), WT or phosphomimetic (S897E) EphA2. We found that cells expressing EphA2(S897E) had lower γH2AX level (**Fig. 5J**), cytoplasmic DNA (**Fig. 5K**) and induction of *Cgas, Sting* and *Ifn* genes (**Suppl. Fig 5A**) when compared to WT EphA2.

While EphA2 inhibition results in DNA damage, the substrate(s) downstream of EphA2 is unclear. Because EphA2 is an RTK, we took a candidate approach by literature search for DNA damage repair (DDR) proteins that are known to be tyrosine-phosphorylated, and these include ATM, RAD51 and JNK.

Specifically, ATM is phosphorylated at Tyr370 by EGFR to promote DNA repair following radiation^23^. Phosphorylation of JNK kinase at Thr183/Tyr185 promotes γH2Ax formation following radiation, leading to apoptotic DNA fragmentation^24^. Bcr-abl fusion kinase phosphorylates RAD51 at Tyr54 and Tyr315, which enhances its recombinase activity to activate RAD51 and promote DNA repair^25^. Of these three proteins, we found that EphA2i treatment resulted in complete loss of phospho-RAD51 (Y315) by western blot (**Fig. 5L**). Conversely, ectopically expressed EphA2 binds and phosphorylates endogenous RAD51 at Tyr315 (**Fig. 5M**), thereby supporting EphA2 as the upstream kinase of RAD51. RAD51 is a recombinase that complexes with single-stranded DNA to enable homology searching and catalyze strand exchange, i.e., homologous recombination following DSBs^26^. In support, we found that RAD51 inhibitor RI-1^27^ caused DNA strand breaks, leakage of DNA into the cytosol and induced expression of *Cgas, Sting* and *Ifn* genes (**Suppl. Fig. 5B, 5C, 5D**). Supporting TAK1 as the upstream kinase of EphA2 and thus RAD51, Takinib decreased both phospho-EphA2(S897) and phospho-RAD51 (Y315) (**Fig. 5N**) Corroborating these results, KPPC cells stably expressing a phosphomimetic EphA2(879E) were more potent in phosphorylating RAD51, and this upregulation was less sensitive to the Takinib, when compared to WT EphA2 (**Fig. 5J**). Furthermore, topoisomerase inhibitor SN38 promoted, whereas Takinib inhibited, the nuclear accumulation of phospho-RAD51 and phospho-EphA2 (**Fig. 5O**). Together, these data support a crucial role of TAK1 in maintaining DNA integrity through phosphorylation of EphA2 and subsequently RAD51, which explains why targeting TAK1 results in DNA damage and activation of the cGAS-STING cascade.

### Takinib reprograms the immune TME and promotes infiltration of activated T cells

To facilitate clinical translation, we tested the therapeutic potential of Takinib in PDAC mice. We treated 6-week-old autochthonous KPPC mice with vehicle or Takinib (50 mg/kg/day by Intraperitoneally) for 2 weeks. Compared to vehicle, Takinib treatment resulted in smaller tumors (**Fig. 6A**), which had lower late PanIN and PDAC lesions, as well as significantly lower collagen (by Sirius Red) and mucin (by Alcian Blue) deposition (**Fig. 6B**). Flow cytometry of Takinib-treated PDAC showed increased total immune, effector CD4^+^, total and effector CD8^+^ T cells, as well as decreased Treg CD4^+^ T cells (**Fig. 6C**).

**Figure 6.**
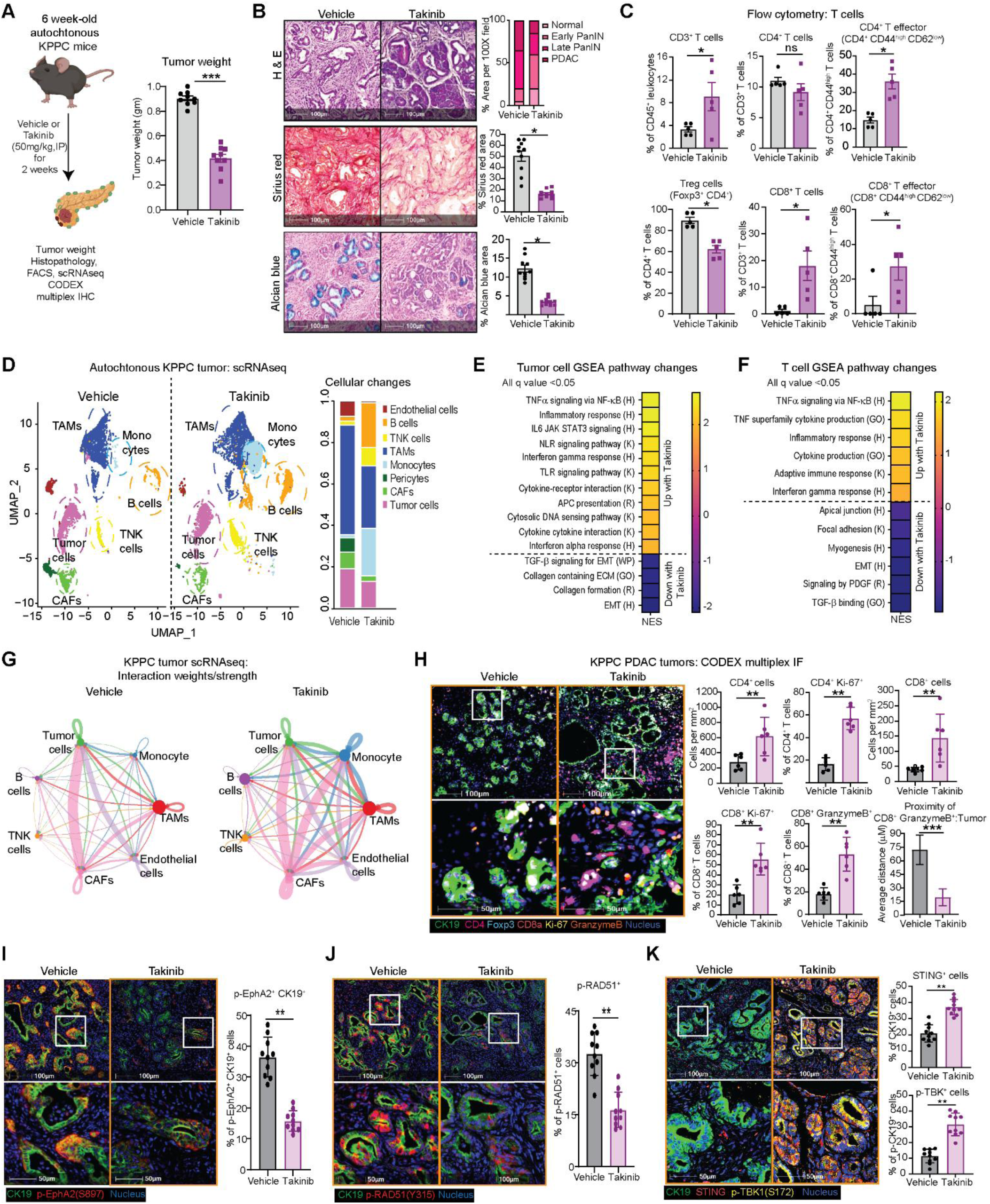
Takinib reprograms the immune TME and promotes infiltration of activated T cells. **(A)** Experimental schematic and final weights of tumors from WT KPPC mice treated with vehicle or Takinib (50mg/kg) IP for 14 days. **(B)** Representative images and quantification of H&E, Sirius red and Alcian blue staining of vehicle and Takinib-treated KPPC tumors. **(C)** FACS analysis of intratumoral indicated T cells (N=6/arm). **(D)** UMAP plot of scRNAseq of vehicle-or Takinib-treated KPPC tumors showing cell subsets (N=4 tumors/arm). Heatmaps showing GSEA signatures in Takinib-treated **(E)** tumor or **(F)** TNK cells based on scRNAseq. Hallmark (H), Gene Ontology (GO), Reactome (R) and Kyoto Encyclopedia for Genes and Genomes (K), database from MSigDB. **(G)** Differential interaction weight/strength between TNK and major cell types from vehicle and Takinib-treated KPPC tumors by CellChat analysis. **(H)** Representative CODEX multiplex IF images and quantification of indicated T cells and proximity analysis. Representative multiplex IHC images and quantification of **(I)** phospho-EphA2^+^ (S897, red) tumor CK19^+^ (green) cells, **(J)** phospho-RAD51 (Y315, red), **(K)** STING^+^ (red) or phospho-TBK^+^ (S172, yellow) tumor CK19^+^ (green) cells. (\**P* < 0.05, \*\**P* < 0.01, \*\*\**P* < 0.001; scale bars 50 and 100 µM).

Additionally, Takinib-treated tumors had decreased number of granulocytes, monocytes, and TAMs (**Suppl. Fig. 6A**), which, by CODEX multiplex IF were skewed towards CD206^-^ (M1-like) polarization (**Suppl. Fig. 6B**). Both TAMs (F4/80^+^) and dendritic cells (CD11c^+^) in Takinib-treated tumors expressed higher levels of MHC II, indicative of enhanced antigen presentation (**Suppl. Fig. 6C**). Interestingly, in Takinib-treated tumors, expression of TGF-β in both CAFs and PDAC cells was significantly lower (**Suppl. Fig. 6D**), supporting our qPCR data that TAK1 controls *Tgfb* gene expression (**Suppl. Fig. 2A**)

To study the systemic impact of Takinib treatment on all cell types, we performed scRNAseq on vehicle- or Takinib-treated KPPC tumors. UMAP clustering revealed an expansion in T and B cells, reduction in TAMs, CAFs and PDAC cells in Takinib-treated tumors (**Fig. 6D, Suppl. Fig. 6E**). GSEA of Takinib-treated PDAC cells showed upregulation of inflammatory signatures including the TNF, TLR, IL-6, IFNα, IFNγ, NOD, cytosolic DNA sensing and antigen presentation, and downregulation of TGFβ, collagen and EMT signatures (**Fig. 6E**). Takinib-treated PDAC cells expressed much higher levels of inflammatory cytokines *Tnf, Il1a, Il1b, Nlrp3* and chemokines *Cxcl2, Cxcl3, Cxcl5, Ccl3* and *Ccl4* (**Supple. Fig. 6F**). These findings are very similar to sh*Map3k7*-silenced KPPC cells (**Fig. 2I, 2J, Supple. Fig. 4A**) and heterozygous *Map3k7*^f/wt^ KPPC mice (**Fig. 3G**) and further validate that TAK1 inhibition results in an inflammatory response in PDAC cells. Focused GESA on T cells showed upregulated cytotoxic T cell signatures including iTNF, inflammatory response and IFNϒ and reduction in focal adhesion, collagen, and ECM signatures (**Fig. 6F**). CellChat analysis nominated increased interaction of TNK and B cells with other cell types (**Fig. 6G**), as well as increased MHC 1 presentation from PDAC cells to TNK cells (**Suppl. Fig. 6G**) in Takinib-treated tumors, which resonated findings in heterozygous *Map3k7*^f/wt^ KPPC tumors. CODEX multiplexed IF showed that Takinib-treated tumors had increased total and proliferating CD4^+^ and CD8^+^ T cells, as well as granzyme B-expressing CD8^+^ T cells that were physically closer to tumor cells (**Fig. 6H**).

As parallel confirmation of our cell line experiments showed that Takinib suppresses phospho-EphA2 and -RAD51 to trigger the cGAS-STING cascade, CK19^+^ PDAC cells in Takinib-treated tumors displayed lower phospho-EphA2 (**Fig. 6I**) and phospho-RAD51 staining (**Fig. 6J**), and higher expression of STING and phospho-TBK1 (**Fig.6K**). Additionally, more PDAC cells in Takinib-treated tumors displayed cleaved-caspase-3, a marker of apoptosis and calcineurin, a marker of immunogenic cell death (**Suppl. Fig. 6H**). Overall, these data support Takinib treatment triggers DNA damage and the cGAS-STING cascade, resulting in immunogenic cell death that triggers adaptive T cell response.

### Genetic or pharmacologic suppression of TAK1 renders immune checkpoint blockade effective

Our data so far led us to hypothesize that suppression of TAK1 promotes adaptive T cells response. To directly prove this, we injected WT and two different heterozygous *Map3k7^f/wt^* KPPC cell lines subcutaneously in C57BL/6J mice. We found that heterozygous *Map3k7^f/wt^* KPPC tumors had much slower growth kinetics (**Fig. 7A**), were smaller at euthanasia (**Fig. 7B**). We also injected WT and heterozygous *Map3k7^f/wt^*KPPC cells orthotopically in Nur77-GFP C57BL/6J mice and found more infiltration of Nur77-GFP^+^ CD8^+^ T cells in *Map3k7^f/wt^* KPPC by FACS (**Fig. 7C**). ScRNAseq analyses of TNK cells from *Map3k7^f/wt^* PDAC tumors showed upregulation of co-stimulatory molecules such as *Cd28, Cd226* and *Cd154*, but also exhaustion markers *Ctla4, Lag3* and *Tigit* (**Suppl. Fig 7A**). We chose to test anti-CTLA4 plus anti-PD1 as these immune checkpoint blockade (ICB) agents are widely used in the clinic. Indeed, while combined anti-CTLA4 and anti-PD1 had no suppressive effect on WT KPPC tumors, it was highly effective in suppressing the tumorigenic growth of *Map3k7^f/wt^* KPPC cells (**Fig. 7D**). We performed similar xenograft experiments using scramble or *shMap3k7*-silenced KPPC cells and observed similar suppressive effect of ICB on *shMap3k7*-silenced KPPC tumors (**Fig. 7E**). In this model, neither anti-CTLA4 nor anti-PD1 alone suppressed the growth of *shMap3k7*-silenced tumors (**Suppl. Fig. 7B**). Furthermore, the suppressive effect of ICB on *shMap3k7*-silenced tumors was completely abrogated when mice were pre-treated with both neutralizing anti-CD4 and anti-CD8 antibodies, but less with either alone (**Fig. 7F, 7G**), suggesting that both CD4^+^ and CD8^+^ T cells are essential for mounting a robust anti-tumor response. Additionally, co-administration of STING inhibitor abrogated effect of ICB in *shMap3k7*-silenced KPPC tumors, supporting that cGAS-STING activation as a critical step in T cell immunity (**Suppl. Fig 7C, 7D**).

**Figure 7.**
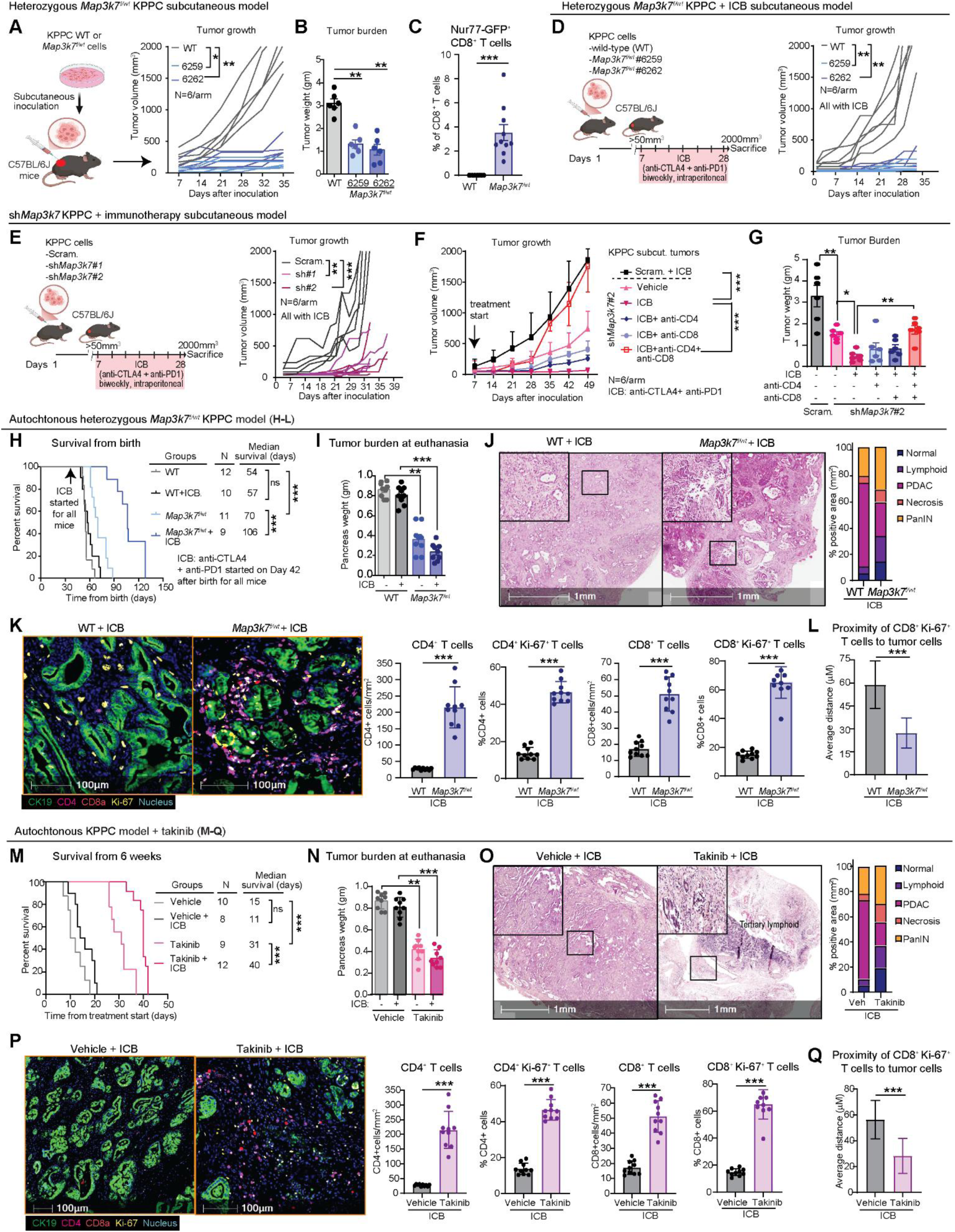
Genetic or pharmacologic suppression of TAK1 renders immune checkpoint blockade effective. **(A)** Experimental schematic, growth kinetics and **(B)** final weight of WT or two different *Map3k7^f/wt^* KPPC tumors in C57BL/6J mice (N=6 mice/arm). **(C)** FACS analysis showing % GFP^+^ CD8^+^ T cells in WT or *Map3k7^f/wt^* KPPC in Nur77-GFP mice. **(D)** Experimental schematics and growth kinetics of **(D)** WT vs M*ap3k7^f/wt^*; or **(E)** scram. vs sh*Map3k7* KPPC cells in C57BL/6J mice treated with dual ICB (anti-PD1 and anti-CTLA4, biweekly, by IP, N=5 or 6 mice/arm). **(F)** Growth kinetics and **(G)** final weight of the indicated scram. or sh*Map3k*7 KPPC tumors in C57BL/6J mice treated as indicated (N=6 mice/arm). WT and *Map3k7^f/wt^* KPPC mice untreated or treated with dual ICB until humane endpoints **(H)** Kaplan-Meier survival plot, **(I)** Tumor burden, Representative images and quantification of **(J)** H&E, **(K)** multiplex IF of T cells and **(L)** Proximity of tumor and CD8^+^ Ki-67^+^ cells. WT KPPC mice treated with ICB with or without Takinib until humane endpoints **(M)** Kaplan-Meier survival plot, **(N)** Tumor burden, Representative images and quantification **(O)** H&E, **(P)** multiplex IF of T cells and **(Q)** Proximity of tumor and CD8^+^ Ki-67^+^ cells (\**P* < 0.05, \*\**P* < 0.01, \*\*\**P* < 0.001; scale bars 1mm or 100 µM).

Next, we tested the effect of ICB on autochthonous KPPC mice, arguably the most challenging PDAC model to treat due to its aggressive and low antigenic tumors^28^. First, we treated WT and heterozygous *Map3k7^f/wt^* KPPC mice with ICB starting from 6 weeks of age, when PDAC has formed^7^, till humane endpoints which include >20% body weight loss from baseline, low mobility and scuffled appearance. Although ICB had no impact on the survival of WT KPPC mice, as expected, it significantly prolonged the survival of heterozygous *Map3k7^f/wt^* KPPC mice (57 vs. 106 days, **Fig. 7H**). At euthanasia, the ICB-treated *Map3k7^f/wt^* tumors were significantly smaller than WT (**Fig. 7I**), and by histology had less PDAC foci and larger areas of necrosis and lymphoid infiltration (**Fig. 7J**). Multiplex IHC showed significantly more total and proliferating CD4^+^ and CD8^+^ in ICB-treated *Map3k7^f/wt^* tumors, as compared to WT (**Fig. 7K**). By proximity analysis, the Ki-67^+^ CD8^+^ T cells in *Map3k7^f/wt^* tumors were closer to PDAC cells (**Fig. 7L**).

To facilitate clinical translation, we tested whether systemic Takinib treatment would recapitulate findings in heterozygous *Map3k7^f/wt^* KPPC. We treated 6-week-old WT KPPC mice with vehicle, Takinib (50 mg/kg/day) alone, ICB alone or combined Takinib and ICB until human endpoints. While Takinib alone prolonged the survival of WT KPPC mice, combined Takinib and ICB led to further prolongation of survival (**Fig. 7M**). At euthanasia, combo-treated tumors were smaller than vehicle or ICB-treated tumors (**Fig. 7N**) and by histologic analyses had much less PDAC foci, more necrosis and lymphoid infiltration (**Fig. 7O**). As seen with ICB-treated heterozygous *Map3k7^f/wt^* KPPC tumors, the combination of Takinib and ICB resulted in higher infiltration of total and proliferating CD4^+^ and CD8^+^ T cells (**Fig. 7P**), as well as closer proximity between proliferating CD8^+^ T and tumor cells (**Fig. 7Q**). Together, we provide genetic and pharmacologic evidence to support combining TAK1 inhibitor with ICB for PDAC patients.

## DISCUSSION

We identify TAK1 as a promising therapeutic target to reprogram the immunologically cold PDAC TME into a more inflamed, “hot” state. Mechanistically, we uncover a previously underappreciated role for TAK1 in maintaining genomic integrity by phosphorylating EphA2 and RAD51. Inhibition of TAK1 or EphA2 leads to DNA strand breaks and subsequent cytoplasmic DNA leakage, which triggers the cGAS-STING cascade and adaptive immune response. Several STING agonists are currently being evaluated in clinical trials as a means to elicit robust adaptive T cell responses^29^. However, their clinical application faces significant challenges, including the risk of excessive immune activation leading to cytokine release syndrome (CRS), and the requirement for intratumoral injection that limits clinical feasibility and systemic efficacy.

We found that TAK1 and EphA2 inhibition results in decreased RAD51 phosphorylation at Tyr315 provide a mechanistic explanation for cGAS-STING activation. Post-translational modifications, including phosphorylation, finely tune RAD51’s activity and stability. Specifically, phosphorylation at Tyr315 enhances RAD51’s recruitment to sites of DNA damage where it nucleates with single stranded DNA, performs homology search and exerts its recombinase activity^30^. We now show that EphA2 is a tyrosine kinase that can bind and phosphorylate RAD51 at Tyr315. In normal cells, EphA2 primarily localized to the plasma membrane, where it interacts with membrane-bound ephrin-A ligands on adjacent cells to regulate cell-cell communication and maintain tissue architecture. However, EphA2 is frequently overexpressed or activated in human cancers. Aberrant expression of EphA2 is associated with T cell exclusion in PDAC. Ablation of EphA2 downregulates expression of prostaglandin endoperoxide synthase 2 (*PTGS2*, which encodes COX-2) and TGF-β signature, leading to infiltration of activated T cells which could be further unleased with ICB^31^. Although the detailed molecular mechanism was unclear, this study at least corroborated our findings that targeting EphA2 resulted in a T cell permissive milieu that could render ICB effective. Our study showed that phosphorylation of EphA2 at S897, at least in part by TAK1, promotes its localization to the nucleus where the RAD51 protein resides and can be phosphorylated by EphA2. Prior to our study, to our best knowledge there has not been any report on the role of TAK1 in the nucleus or its direct role in DNA repair. Although we excluded ATM and JNK as substrates for EphA2, other nuclear substrates involved in DDR and mediated by EphA2 remain possible. In summary, we uncover TAK1 as a multifaceted regulator of immune evasion and genomic integrity in PDAC. By elucidating a novel TAK1-EphA2-RAD51 axis and its role in maintaining genomic integrity and suppressing cGAS-STING-mediated immunity, our study provides mechanistic insight and a strong rationale for therapeutic targeting of TAK1 in combination with ICB.

## ACKNOWLEDGEMENTS

Funding support: NIH R37CA219697, R01CA262414, P50CA196510 and The Foundation for Barnes-Jewish Hospital.

