## Supplementary figures and images for "Targeting tumor-intrinsic TAK1 triggers anti-tumor immunity and sensitizes pancreatic cancer to checkpoint blockade"

### Supplementary Figure 1

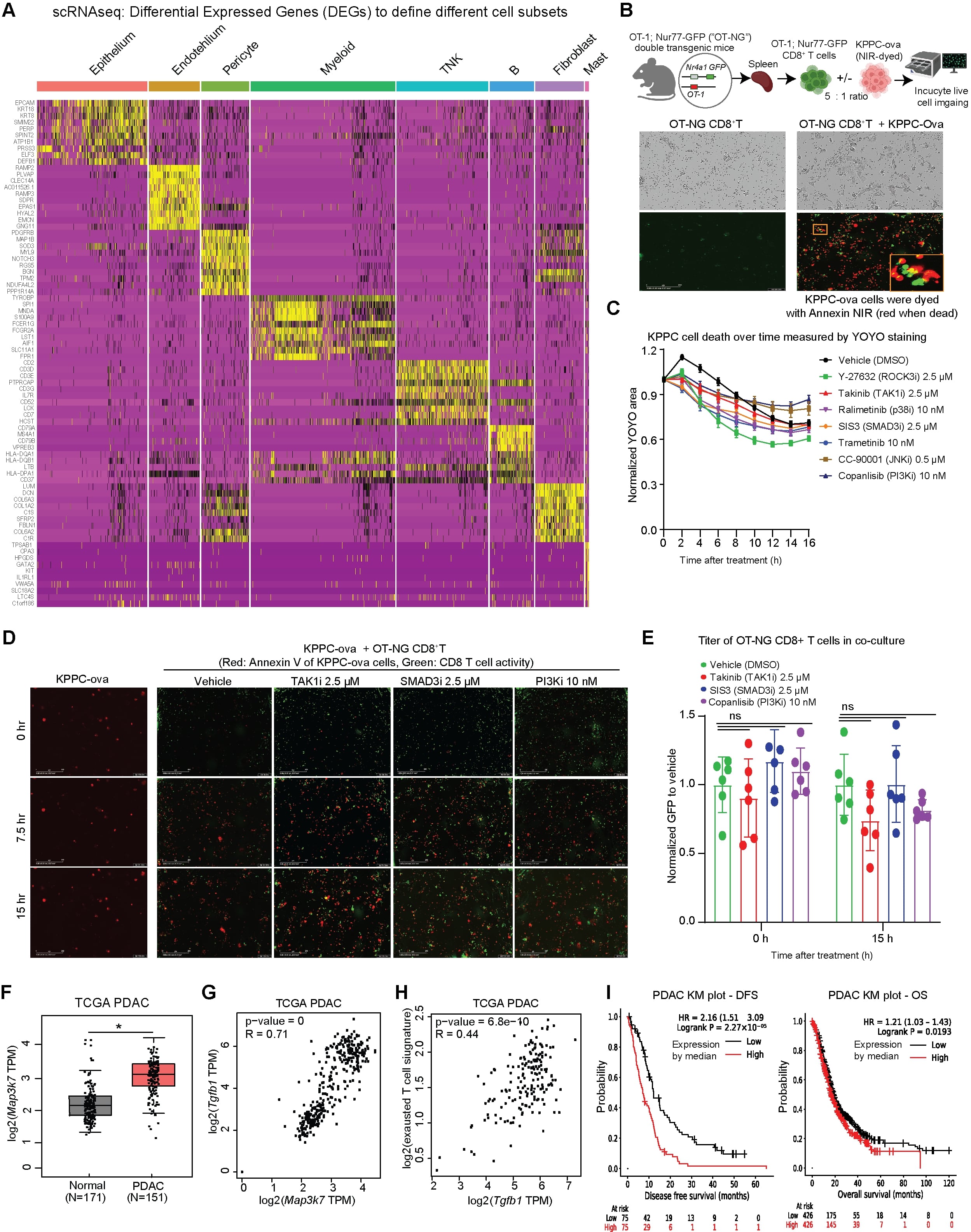

### Supplementary Figure 2

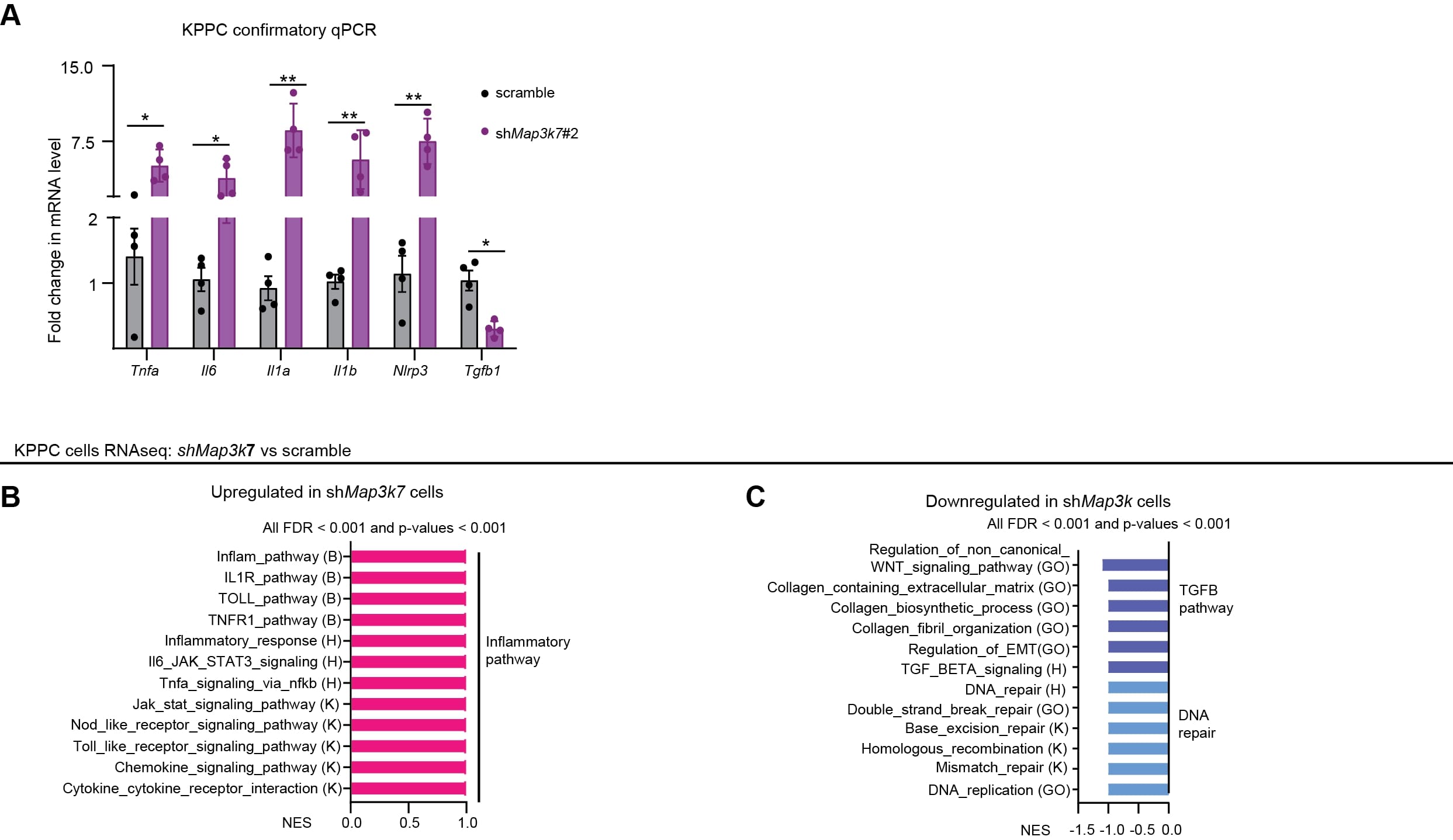

### Supplementary Figure 3

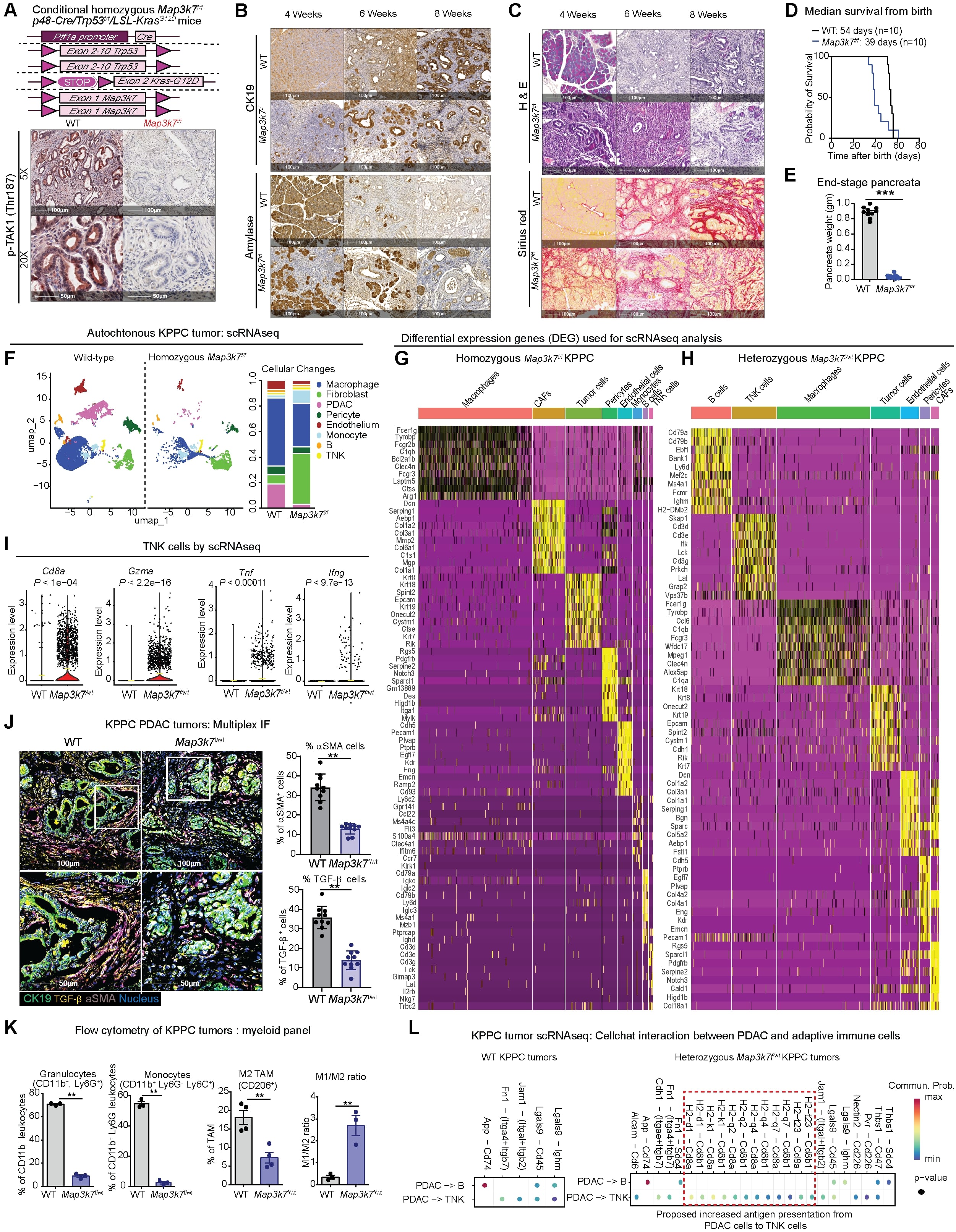

### Supplementary Figure 4

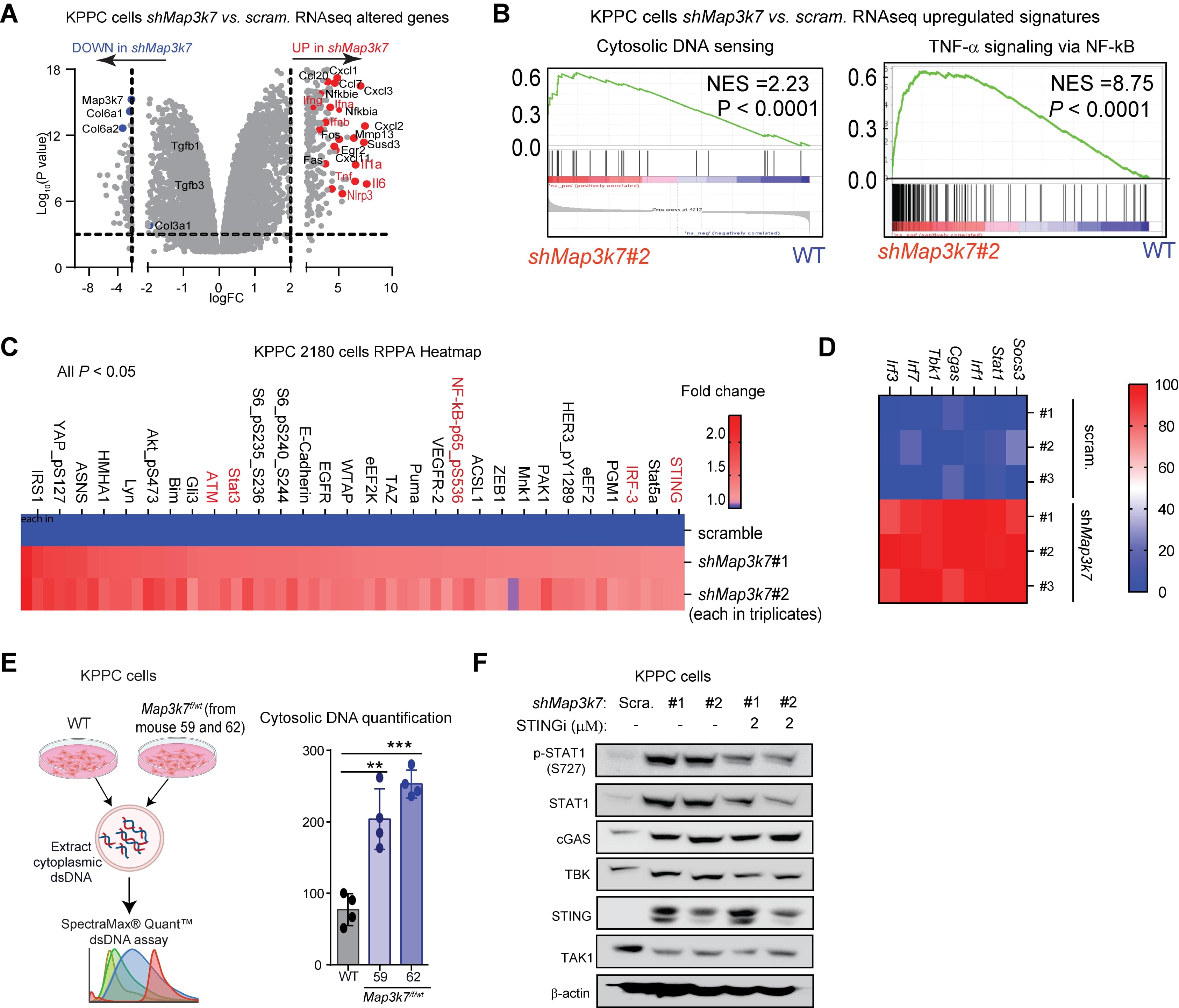

### Supplementary Figure 5

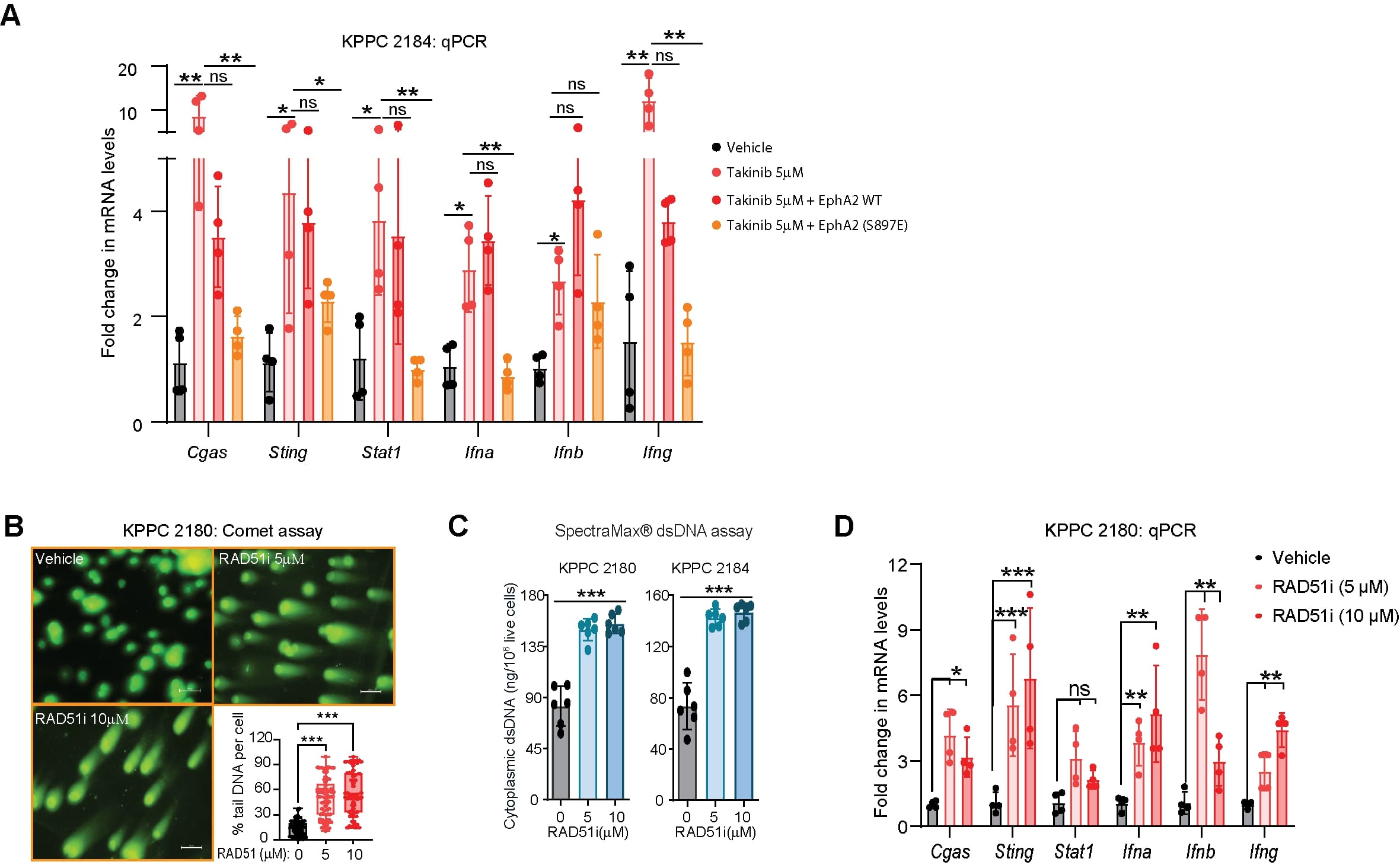

### Supplementary Figure 6

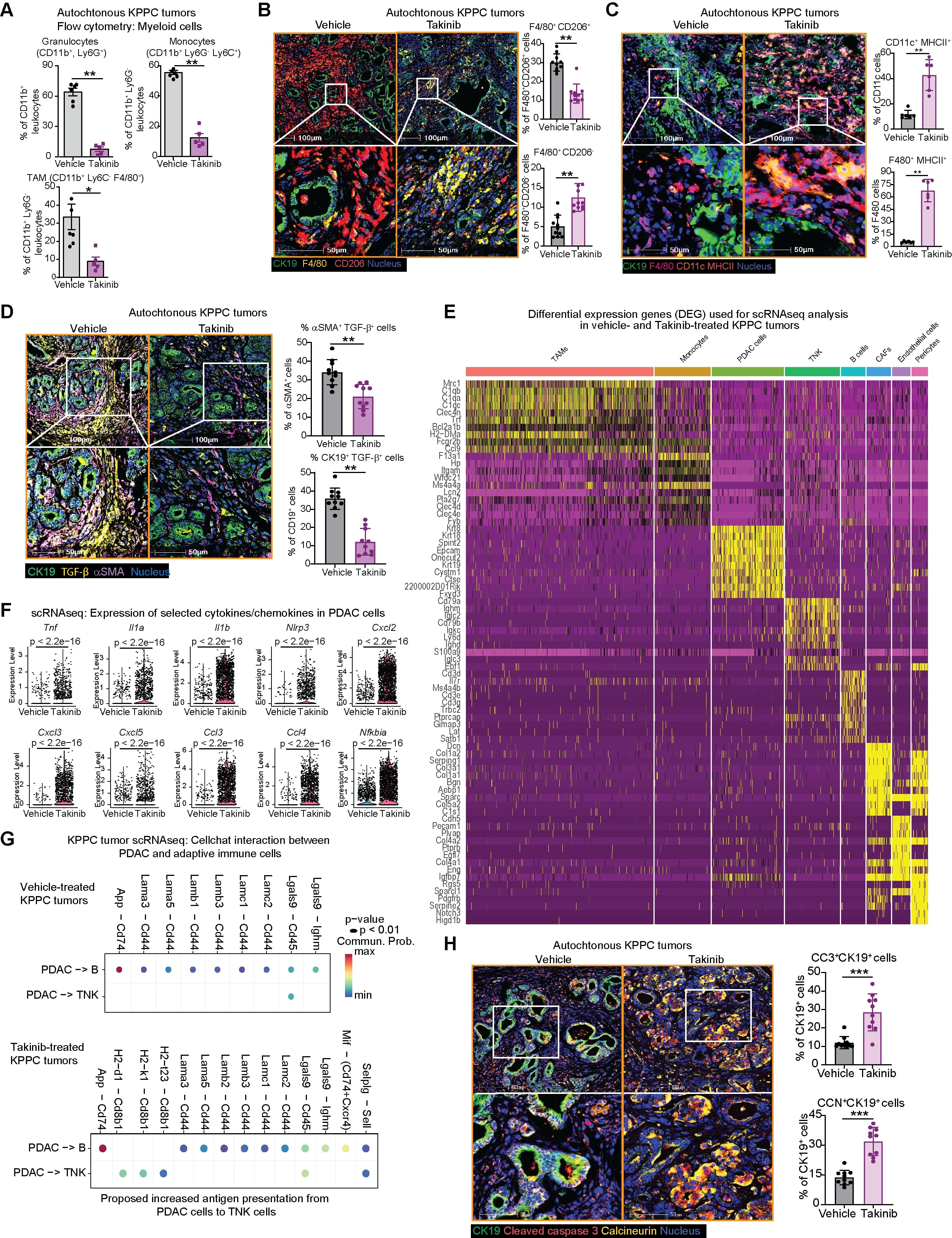

### Supplementary Figure 7

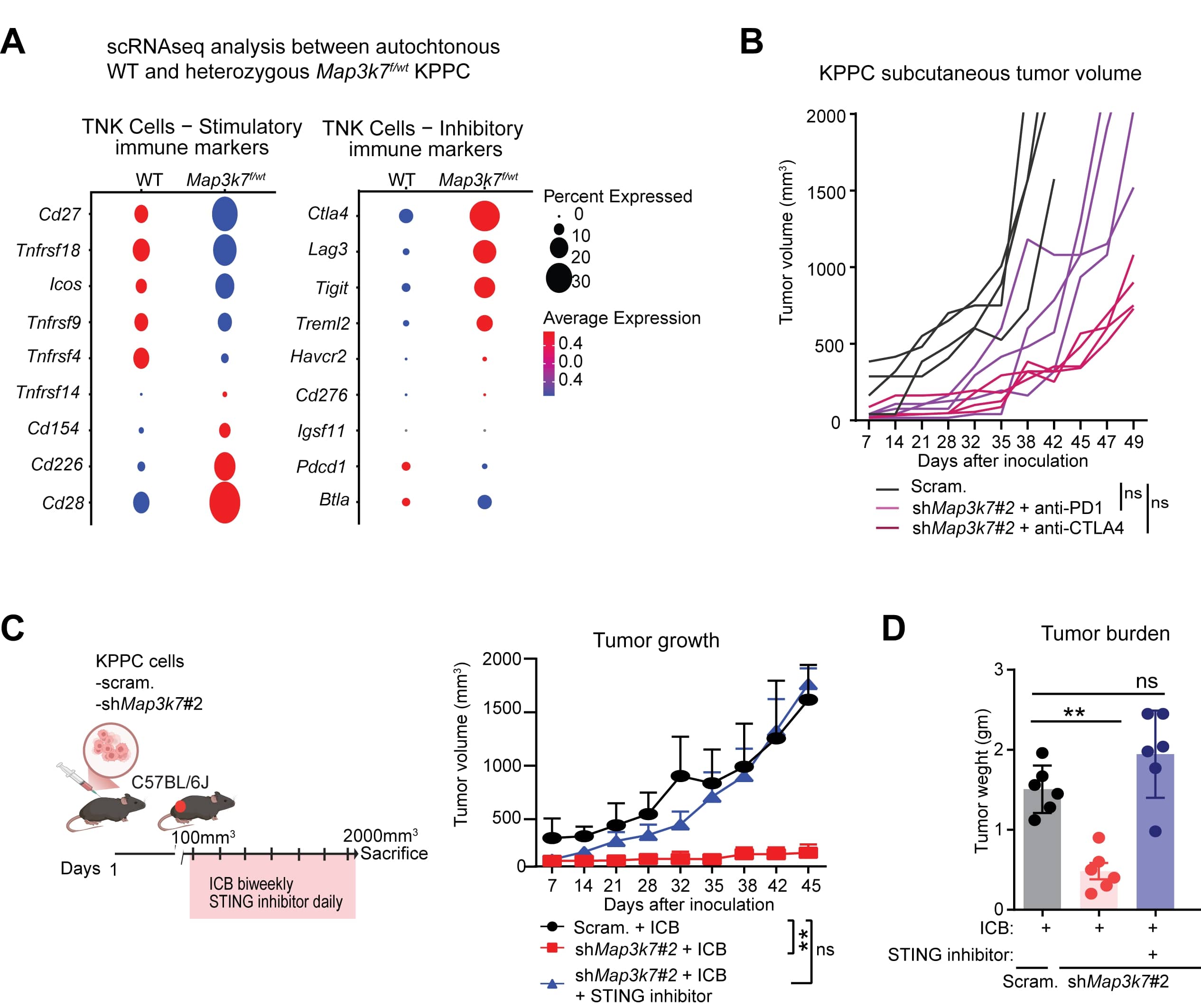
